# Neural tracking as a diagnostic tool to assess the auditory pathway

**DOI:** 10.1101/2021.11.26.470129

**Authors:** Marlies Gillis, Jana Van Canneyt, Tom Francart, Jonas Vanthornhout

## Abstract

When a person listens to sound, the brain time-locks to specific aspects of the sound. This is called neural tracking and it can be investigated by analysing neural responses (e.g., measured by electroencephalography) to continuous natural speech. Measures of neural tracking allow for an objective investigation of a range of auditory and linguistic processes in the brain during natural speech perception. This approach is more ecologically valid than traditional auditory evoked responses and has great potential for research and clinical applications. This article reviews the neural tracking framework and highlights three prominent examples of neural tracking analyses: neural tracking of the fundamental frequency of the voice (f0), the speech envelope and linguistic features. Each of these analyses provides a unique point of view into the human brain’s hierarchical stages of speech processing. F0-tracking assesses the encoding of fine temporal information in the early stages of the auditory pathway, i.e., from the auditory periphery up to early processing in the primary auditory cortex. Envelope tracking reflects bottom-up and top-down speech-related processes in the auditory cortex and is likely necessary but not sufficient for speech intelligibility. Linguistic feature tracking (e.g. word or phoneme surprisal) relates to neural processes more directly related to speech intelligibility. Together these analyses form a multi-faceted objective assessment of an individual’s auditory and linguistic processing.

## 1. Introduction

Understanding speech is a complex process that relies on activation and cooperation between various brain regions. Different characteristics of incoming speech are processed in different brain regions. Roughly, purely acoustic processing of the speech occurs in subcortical areas and the primary auditory cortex. In contrast, segmentation of words and phonemes occurs in temporal regions of the brain, and integration of words into their context occurs in language-related brain regions, such as superior temporal gyrus and inferior frontal gyrus (Brodbeck et al., 2018c,a). However, only if all stages in this neural pathway are successful speech understanding can be achieved.

Audiologists rely on an extensive test battery to assess a person’s speech understanding. A commonly performed test is to let a subject recall a list of sentences. The outcome of this test expresses speech understanding as a percentage of correctly recalled words. However, such behavioural tests have some disadvantages. First, the subject must listen actively to the stimulus and recall the words. Although this seems like an easy task, it can be challenging or impossible for many populations: persons with locked-in syndrome, young children, persons with aphasia, etc. Although they might understand the speech, they might not be able to recall the heard words. Second, the outcomes of these tests do not pinpoint the origin of the deficit. Is the deficit situated cortically, indicating an issue with the higher-order language processing, or peripherally, suggesting a hearing loss? Third, these behavioural tests rely on highly controlled standalone sentences or words spoken by a professional speaker. Such speech material has limited contextual information. Therefore, it does not resemble a typical day-to-day listening environment.

One can use a more objective approach, such as neurophysiological measures, i.e., metrics derived from brain signals to overcome these issues. Traditional neurophysiological measures, like the auditory brainstem response (ABR), the auditory steady-state response (ASSR) or the frequency following response (FFR), require EEG measurement. During such a measurement, a participant listens to repetitive presentations of a short sound stimulus (for a review, see Picton (2010)). Typical stimuli include clicks, tones, chirps and vowels. The repetitive stimulation is necessary as response instances need to be averaged to reduce measurement noise, but it is highly unnatural and demotivating for the listener (Theunissen et al., 2000; Hamilton and Huth, 2018). In recent years, technical advances in data analysis have made it possible to analyse neural responses measured while a participant listens to continuous natural speech without repetition (for a review, see Brodbeck and Simon, 2020). These neural responses time-lock to the presented speech and this phenomenon is called neural tracking. Measuring neural responses to continuous natural speech was originally proposed by Lalor et al. (Lalor et al., 2009; Lalor and Foxe, 2010) and the methods were further developed by, amongst others, Ding and Simon (2012a,b), O’Sullivan et al. (2015) and Crosse et al. (2016a).

The possibility of investigating continuous speech processing by measuring neural tracking is an important innovation. Humans do not communicate with the stimuli of traditional objective measures: repetitive tones or clicks. Context-rich continuous speech better approximates natural language use, and as a result, research findings with these stimuli are more relevant for auditory processing in day-to-day communication (Kei et al., 1999; Pichora-Fuller et al., 2016; Hamilton and Huth, 2018; Keidser et al., 2020). Moreover, continuous speech is more comfortable and enjoyable for the listener. The stimulus can even be targeted towards the population of interest: e.g. a fairy tale for young children or a podcast for adults. When participants are interested in the content of the stimulus, they maintain attention for longer, and as a result, the neural response measurement may be of higher quality. Finally, natural speech stimuli are better suited for research with hearing aids. Hearing aid signal processing is designed specifically for natural speech and may behave unpredictably with artificial sounds, corrupting the experiment.

In this article, we will give an overview of neural tracking of continuous speech with primary emphasis on neural tracking of single-talker speech. However, a promising and emerging related field is the use of neural tracking to determine the focus of attention, called auditory attention decoding (AAD). When listening to a target speaker in a mixture of multiple speakers, higher neural tracking is observed for the attended speaker than ignored speakers. Auditory attention decoding may enable smart hearing devices that can reinforce the attended speaker while attenuating the unattended speakers (Geirnaert et al., 2021).

In this article, we evaluated neural tracking as a diagnostic tool to assess multiple levels of the auditory pathway. Such a tool would be based on one EEG recording of a person listening to natural speech, which is analysed in various ways to assess whether there is a speech understanding deficit and if so, where it is situated in the auditory pathway (e.g., peripherally or cortically).

## 2. Methods to measure neural tracking

The most common approach for measuring neural tracking are linear, decoding or encoding models. These models measure the amount of neural tracking, i.e., how strongly the neural responses time-lock to a stimulus feature and are discussed in more detail in the following subsections.

Measuring neural tracking requires two inputs: neural responses in the form of single-channel or multi-channel EEG (or MEG) and one or more features representing the stimulus (see section 3). A linear relation between the EEG and the stimulus feature is modelled to investigate how well the stimulus information is encoded in the neural activity. Linear modelling is possible in two ways: reconstructing the feature from the EEG (decoding, section 2.1) and, conversely, predicting the EEG from the feature (encoding, section 2.2) or a combined approach (CCA (de Cheveigné et al., 2018) as discussed in section 2.5). As discussed below, the decoding and encoding analyses provide different but complementary information about neural tracking.

### 2.1 Decoding modelling

In decoding modelling, one reconstructs the stimulus feature from a weighted sum of the EEG signals of the different channels and their time-shifted versions. The time-shifted versions are included to account for the time difference between the sound and the associated neural response.

The decoding modelling procedure is visualised in panel A of Figure 1. First, the weights that provide the optimal reconstruction are determined based on the time-shifted EEG and its corresponding stimulus feature. Then those weights are applied to the EEG, resulting in a reconstructed stimulus feature. The reconstructed feature is correlated with the actual stimulus feature of the test data to determine the reconstruction accuracy. This reconstruction accuracy is a measure of neural tracking as it indicates how well the stimulus information is time-locked with the EEG. A higher reconstruction accuracy will therefore reflect higher neural tracking of the speech.

**Figure 1:**
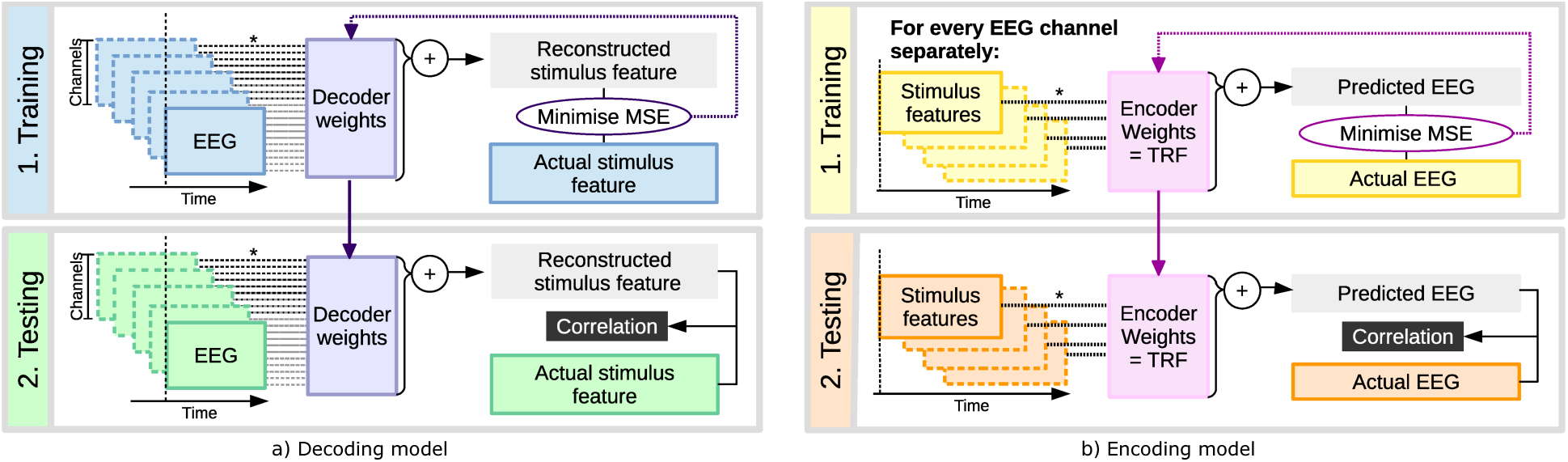
*A. Schematic representation of decoding modelling.* The stimulus feature is reconstructed in decoding modelling based on a linear combination of time-shifted EEG data. In the training phase, the model is estimated by optimising the decoder weights to minimise the MSE (mean squared error) between the reconstructed stimulus feature and the actual stimulus feature for a training data set. Then, the weights are applied in the testing phase to reconstruct the stimulus feature for the testing dataset. The final output is the correlation between the reconstructed and actual stimulus features for the testing dataset. The division in training and testing dataset is done according to the cross-validation technique (described in 2.4) *B. Schematic representation of encoding modelling*. In encoding modelling, the EEG data in each EEG channel is predicted based on a linear combination of time-shifted stimulus features. Again, the encoder weights or TRFs (temporal response functions) are estimated by minimising the reconstruction MSE for a training data set. Then the TRFs can be studied as is or used to predict the EEG for a testing data set. The output of the testing phase is the correlation between the predicted EEG and the actual EEG. The division in training and testing dataset is done according to the cross-validation technique (described in 2.4)

The decoding modelling approach is a powerful analysis tool since the information of multiple EEG channels (often 32 or more) can be combined to reconstruct a stimulus feature with often only one dimension (although multi-dimensional features are possible).

A disadvantage of decoding modelling is that the weights of the model are extraction filters which cannot and should not be interpreted to investigate the spatial pattern of the response (Haufe et al., 2014). Extraction filters do not always have large weights when the corresponding EEG channels contain a lot of response information. When an EEG channel captures information about a noise component, it can be used in the modelling process to remove the noise component from other EEG channels. As a result, some channels may receive large weights because they are helpful for noise reduction purposes and not because they contain response information (Montoya-Martínez et al., 2021). Haufe et al. (2014) defines an inversion method to make the topography of the decoding model interpretable, or one can use a encoding model.

Another disadvantage of decoding modelling is that the evaluation of the model, i.e., the loss function, should be adapted when investigating the reconstruction accuracy of sparse, impulsive features. These sparse, impulsive features can, for example, code the onset of a phoneme or word. They consist of an array of zeroes with a given value at the onset of a word or phoneme. Such features cannot be reconstructed from the continuous EEG signals with a linear model. Therefore, if sparse, impulsive features are used, a correlation between the actual and reconstructed features is sub-optimal. The evaluation of a decoding model, i.e., taking a correlation, should be adapted when using sparse, impulsive features. Another option is to equalise the feature spectrum to the EEG spectrum by convolving the sparse, impulsive feature with an appropriately smooth kernel. However, these approaches remain to be investigated in detail, and typically encoding models are used with impulsive features.

### 2.2. Encoding modelling

Encoding modelling can be used to study the spatio-temporal properties of the response: the EEG signal in each channel is predicted from the speech features via a weighted sum of the different speech features and their timeshifted versions. These weights form a temporal response function (TRF) for each speech feature and each EEG channel. A TRF consists of the estimated weights at the different time-shifts of the speech feature, reflecting how the EEG response is modulated by the stimulus at different time-shifts. The encoding model can deal to some extent with autocorrelation of the speech stimulus. Autocorrelation denotes that speech is correlated with itself at different time lags. Because the computation of the encoding model uses this autocorrelation, it can prevents the smearing over time of the TRF (for more details, see Crosse et al., 2016b).

Panel B of figure 1 schematically presents the encoding modelling process. Note that for the encoding modelling, the time-shifting occurs in the opposite direction than for decoding modelling. Each EEG channel is considered separately, so encoding models cannot reduce noise in the EEG signal by combining information across channels. The advantage of this approach is that the TRF weights are activation patterns and not extraction filters. Activation patterns have large weights for the EEG channels containing a lot of response information and can therefore be interpreted. The temporal aspects of these TRF weights are particularly interesting to investigate the neural response latency. Depending on the considered brain area, different bottom-up neural response latencies can be expected: about 5-10 ms for auditory processing in the upper brainstem and at least 12-30 ms for processes in the primary auditory cortex (Tichko and Skoe, 2017; Brugge et al., 2009). Higher-order cortical processes that modulate the neural response, like attention and interpretation of the speech, occur with delays of 200 ms or more (for a review, see Martin et al., 2008). Similarly to decoding modelling, a measure of neural tracking can be extracted by correlating the actual measured EEG responses with the predicted EEG responses. This correlation is called the prediction accuracy and is a measure of neural tracking, i.e., the better the brain tracks the speech, the higher the prediction accuracy.

Compared to decoding modelling, encoding modelling results in a lower magnitude of neural tracking. The reasons are twofold. Firstly, in encoding modelling, the actual EEG is correlated with the EEG predicted from the speech features. However, the predicted EEG is a gross simplification of the content of the actual EEG signals, which contain responses to the speech together with a plethora of non-speech-related EEG activity and noise. Secondly, encoding modelling cannot use across-channel information to reduce noise in the EEG signals (Das et al., 2019). However, a lower magnitude of neural tracking does not necessarily mean that the encoding model is less valid or reliable than the decoding model. The only issue is that the metric to assess the quality of the model is noisy and has lower values.

For each channel, the TRF can be interpreted as the impulse response of the measured auditory system: the information in the input stimulus, i.e., the speech feature, is transformed with this impulse response to produce the output response, i.e., the preprocessed EEG. Please note that the impulse response depends on the preprocessing (further described in section 2.4). The channel-specific TRFs are noisy and therefore often averaged over a selection of EEG channels and subjects. Based on the time-shifts that receive large weights for many of the EEG channels/subjects, the dominant latencies of the response can be derived. These latencies (or delays) can then be used to estimate which stages of neural processing along the auditory pathway contribute to the response. The spatial properties of the response can be further investigated from the distribution of the magnitude of TRF weights over the scalp. This information is usually visualised on a topographic map. Examples of TRFs and topographic maps are shown in figure 4, which is discussed below. The TRFs and the corresponding topographic maps are similar to ERPs with the advantage that they can be computed using a continuous signal. Moreover, the prediction accuracies of the encoding model can also be visualised on a topographic map. Note that such topographic maps only divulge spatial information on scalp level, where the electrodes were located. To study the actual sources of the neural responses within the head, the information form electrode space should be transformed to neural source space (e.g. Brodbeck et al., 2018c).

Although the encoding modelling approach can deal to some extent with the autocorrelation of the stimulus, it can still be problematic for certain features, like the fundamental frequency (further discussed in section 4).

### 2.3. Algorithms to calculate decoding models and encoding models

Different algorithms exist to acquire decoding and encoding models. In essence, the algorithms have the same goal: minimizing the error between the reconstructed or predicted signal with the actual signal. However, due to the noisy nature of EEG signals and the fact that only a small fraction of the EEG signal is auditory stimulus related, this question is ill-posed meaning that multiple solutions are possible. Therefore, these algorithms may yield slightly different outcomes as they might rely on different priors. In this overview, we will focus on two algorithms: ridge regression (Machens et al., 2004) and boosting (David et al., 2007). Both algorithms are supported by a dedicated toolbox, respectively the mTRF toolbox (Crosse et al., 2016b) and Eelbrain (Brodbeck et al., 2021b).

The ridge regression algorithm minimises the mean-squared error between the predicted or reconstructed signal and the actual signal. It relies on the inverse of the autocorrelation matrix of the time-shifted input. The input depends on the considered model, i.e. EEG for a decoding model and speech features for the encoding model. Taking the inverse of this autocorrelation matrix is ill-posed as the rows of the matrix are mutually dependent, because the time-shifted inputs are dependent. To solve this, a ridge parameter is added to each diagonal element of the autocorrelation matrix. Using a cross-validation approach (described in 2.4), the ridge parameter can be determined. To do so, the measure of neural tracking is calculated for multiple values of the ridge parameter (for example ranging from 10^−2^ to 10^3^ in steps of the powers of 10). The ridge parameter with the highest accuracy is selected and used for the analysis.

Like ridge regression, the boosting algorithm minimises the error between the predicted or reconstructed signal and the actual signal, and aims for a maximally sparse solution. In contrast to ridge regression, which has a closed-form solution, boosting relies on an iterative approach to determine the model’s weights. Initially, all weights are set to zero. Subsequently, each weight is changed by a specific value. The error between the predicted or reconstructed signal and the actual signal is calculated for every change. The weight which resulted in the smallest error is selected. Then, these weights are used to start the next iteration. These iterations are performed until a stopping criterion is met, e.g. the error stops decreasing, or the number of iterations exceeds the limit.

Although both algorithms lead to a similar results, the outcome has some apparent differences (Kulasingham and Simon, 2022). Ridge regression results in smooth solutions which are more spread across time and channels while boosting results in a more sparse solution which is better defined in time and space. On the other hand, as boosting relies on this iterative approach, it is more computationally expensive than ridge regression.

### 2.4. Preprocessing and model evaluation

Preprocessing of the EEG and the speech features affects the results and interpretation of the model. Especially the interpretation of the patterns in the TRF should be made carefully with respect to the preprocessing characteristics.

The filtering method is a crucial aspect to consider when preprocessing the EEG data. Every filter has specific characteristics which affect the impulse response of the system. Two important filter characteristics should always be considered: the causality and the filter’s phase response. The causality relates to which data points the filter uses. For causal filters, only past data points can affect the output of a specific data point, while for acausal filters, the output can be affected by past and future data points. The phase of the filter is also important as it denotes the delay introduced by the filter.

Especially when interpreting the TRF, the filter characteristics should be considered. Firstly, we want to emphasise that using a causal filter makes more sense: the stimulus evokes a response in the brain, so only past data points can influence the output. However, a causal filter cannot be zero-phase, and therefore, introduces a delay which should be accounted for. An exception is when the EEG and stimulus features are filtered the same way; the delay will affect both in the same way, and thus the time delay and time-locking between EEG and stimulus is preserved. It should be emphasised that all filter characteristics should reported for a study, as pointed out by de Cheveigné and Nelken (2019). The best practice is to filter the stimulus and the EEG similarly. Additionally, the EEG and stimulus spectrum will become similar, leading to higher prediction or reconstruction accuracies. Regarding decoding modelling, the filter causality and phase are less critical as the weights of the decoding model are not interpreted.

Another critical preprocessing step is the referencing of the EEG signals. Different choices can be made: the central electrode Cz, mastoids or a common average. The reference choice does not affect the overall magnitude of the reconstruction or prediction accuracy, but it does affect the weights of the model. This aspect is not essential for decoding models as the weights cannot be interpreted. For encoding models, the TRF and the distribution of prediction accuracies across the EEG channels are affected by the reference choice. If the EEG signals are referenced to one or a combination of couple of electrodes such as the two mastoids, the obtained referenced EEG favours specific brain regions. If this is not wanted, the common average referencing is a good choice which allows a broader picture of the neural activity. However, when making claims about neural activity in dedicated regions in the brain, using source localisation techniques is a better solution than deriving conclusions based on the TRF.

Preprocessing of EEG also incorporates artefact removal. The above-discussed linear models investigate the timelocked relationship between stimulus features and the EEG responses. As artefacts due to eye blinks or movement are not strictly time-locked to the speech, the models can cope with these artefacts if sufficient data is provided. Nevertheless, artefact suppression should always be considered. Multiple options are possible: multi-channel Wiener filtering (Somers et al., 2018), independent component analysis, denoising source separation (Särelä et al., 2005), multiway canonical correlation analysis (de Cheveigné et al., 2019), etc. These techniques are useful to suppress various artefacts in the EEG signal.

After preprocessing the data, the linear models can be estimated. We want to emphasise the necessity of crossvalidation to estimate and evaluate the models. If the model is estimated and evaluated on the same data, the results are likely to be biased: inflated reconstruction or prediction accuracies and distorted TRF patterns. For example, more peaks might be seen in the pattern because the model learnt the noise in the data. Altogether, estimation and evaluation of the model on the same data leads to unreliable results specific to the used data and may not generalise well to new, unseen data. This can be avoided by using the cross-validation technique (Crosse et al., 2021). This technique relies on a training set, i.e., part of the data used for model estimation and a testing set, i.e., another part of the data, unseen during the model estimation, to evaluate the model on (visualised in Figure 1). The cross-validation technique divides the data into *n* folds of equal length. Subsequently, the model is estimated on *n* − 1 folds and evaluated on the left-out fold. This is repeated until all folds have been used to evaluate the model. The TRFs and prediction or reconstruction accuracies obtained by respectively model estimation and evaluation are then averaged across the different folds. This technique allows identifying a robust model that generalises to new, unseen data.

### 2.5. Other methods to extract neural tracking

Decoding and encoding models imply a directionality: either the stimulus feature is reconstructed or the EEG responses are predicted. Another linear approach is canonical component analysis (CCA), which operates bidirectionally. CCA transforms both EEG and stimulus, so they are maximally correlated, thereby combining the advantages of the decoding and encoding models. Stimulus dimensions irrelevant for measurable responses are removed, as are EEG dimensions irrelevant for auditory perception. Although CCA is a flexible tool that can discover more complex relations than a simple encoding or decoding model, it has more parameters and with this a higher risk of overfitting. Therefore, an appropriate cross-validation strategy is needed, or one has to use dimensionality reduction or regularisation. Furthermore, as with all techniques based on least-squared minimisation, it is prone to outliers. CCA decomposes the signals into multiple components. Although this can ease the interpretation of the results, it should be done with care. CCA orders the components based on their correlation, however a high correlation does not guarantee a physiological interpretation. For example, components that have lowpass or narrowband filtering can have very high correlations. Finally, there is no exact match between the components and the actual neural sources which can complicate the interpretation of the underlying neural basis of the component. This technique has been used in de Cheveigné et al. (2018) and O’Sullivan et al. (2021)

Another linear analysis method is a cross-correlation (Kong et al., 2014; Aiken and Picton, 2008; Petersen et al., 2016; Aljarboa et al., 2022). Here, the cross-correlation is computed between the EEG channels and the speech features. The cross-correlation is computationally inexpensive and can give some insight into the neural responses. Similar to the encoding model, it cannot integrate multiple channels. An disadvantage compared to encoding and decoding modelling, is that the cross-correlation is more sensitive to the autocorrelation of the stimulus, leading to patterns that are smeared out over time. Another disadvantage is that cross-correlation cannot be applied on an unseen test dataset. A comparison of the TRF to the cross-correlation is nicely shown in Crosse et al. (2016b).

Non-linear techniques have been explored to overcome the inherent limitations of linear models. Mutual information is a metric that uses information theory, which captures the shared information between the EEG responses and the stimulus expressed in the unit ‘bits’. Therefore, higher mutual information indicates higher neural tracking. Because it does not make explicit assumptions about the relationship between the stimulus and EEG, it can capture nonlinear aspects. Moreover, mutual information can be calculated between the EEG and the time-lagged stimulus to understand how the mutual information metric behaves for multiple time lags. This results in a pattern similar to the TRF. However, an important consideration is that autocorrelation of the stimulus is not taken into account. Note that the mutual information method does not need to be applied between stimulus and EEG necessarily, but can also be applied between different EEG signals to get additional insights. This technique has been used by Gross et al. (2014), Zan et al. (2020), and Kaufeld et al. (2020).

Another measure of neural tracking can be obtained with a match-mismatch paradigm. In this case, a model is trained to classify whether a given EEG segment is matched or not with a given stimulus segment. This can be done for a single stimulus segment or N segments, in which case the model classifies which of the N segments is matched with the EEG. In the case of decoding or encoding linear models, the stream with the highest prediction or reconstruction accuracy can be identified as the matched stream. The accuracy, i.e., the percentage of correctly identified matched speech streams, is a metric of neural tracking. Please note that auditory attention decoding (AAD) relies on the same principle. Instead of presenting a match and mismatch speech stream, the attended and unattended speech streams are given as input to the model (e.g. Fuglsang et al., 2020; Das et al., 2018; Deckers et al., 2018; O’Sullivan et al., 2015; De Cheveigné et al., 2021).

This paradigm can also be solved in a non-linear fashion with neural networks (e.g. Accou et al., 2021; Monesi et al., 2021; Bollens et al., 2022). Accou et al. (2021) showed that the accuracy of a neural network solving a match-mismatch task could be used to estimate the speech reception threshold. Therefore, this neural network allows for an evaluation of speech understanding based on the EEG responses. Although neural networks can model nonlinear relationships between the EEG and stimulus, they give limited insight into how the network handles the EEG responses. Therefore, evaluating different levels of the auditory system becomes more challenging as it is difficult to tell which information and how the neural network uses it.

Although other methods can quantify neural tracking, in the continuation of this overview, we focus on decoding and encoding models.

## 3. The stimulus feature

The stimulus feature is derived from the presented speech and reflects how a particular speech characteristic varies over time. Many stimulus features can be used, ranging from low-level acoustic characteristics (e.g. the acoustic envelope) to high-level linguistic information (e.g. word surprisal). This flexibility makes the neural tracking framework highly versatile and allows for evaluating multiple levels of the auditory system. It also underlies one of the most prominent advantages of the framework: a single EEG measurement can be analysed with various features of the stimulus and provides information on a range of auditory/language processes. Note that the feature choice is arbitrary, and thus different features will reflect the different stages of the auditory pathway. In this manuscript, we focus on f0 tracking, envelope tracking and linguistic tracking to target different stages in the auditory pathway. Other features are possible and have been investigated in other studies, e.g. word category (Brennan and Hale, 2019), acoustic onsets (Brodbeck et al., 2018a), the spectrogram (Di Liberto et al., 2015), phonetic features (Di Liberto et al., 2015), etc.

In the following sections, we will illustrate the use of neural tracking measures using three prominent (groups of) stimulus features corresponding to three types of neural tracking analyses. We discuss these following the hierarchical organisation of the auditory pathway: starting with auditory processing of the fundamental frequency (f0, section 4), which happens mainly in the subcortical stages of the auditory pathway, then moving on to envelope processing (section 5) which happens in the auditory cortex and ending with linguistic processing (section 6) which happens in the language network of the brain. We focus on how these stimulus features can be used to investigate different aspects of speech processing and different parts of the auditory pathway. Moreover, we provide example results and review findings from relevant studies, including how the responses relate to important clinical measures like hearing thresholds and speech perception. Please note that it is out of the scope of this paper to comprehensively review all stimulus features and related analyses that have been published.

### 3.1. Inter-feature correlation and feature evaluation

All the speech features are derived from the same speech stimulus. This leads to a high inter-feature correlation. For example, at every impulse of a linguistic speech feature, there is a word or phoneme onset, which tends to be associated with a high burst of acoustic energy. In Figure 3, we visualised the inter-feature correlation (panel A). Moreover, we created a TRF to predict the envelope based on the remaining stimulus features (using the boosting algorithm; panel B). Not surprisingly, the envelope feature can be predicted from the other speech features. Even some aspects of linguistic features (such as word surprisal) explain some variance in the envelope feature. This inter-feature correlation can affect the model performance and complicate the interpretation of the results.

This inter-feature correlation can bias the interpretation of results, which is important to consider when investigating the tracking of a linguistic feature using a model with only the linguistic feature of interest. When significant neural tracking is observed, it cannot be attributed to solely the brain tracking linguistic aspects of speech due to this interfeature correlation (Daube et al., 2019).

To overcome this issue, a good approach is to assess the feature’s contribution to the model performance. To do so, the features of interest must be defined together with control features, i.e., features that are not of interest in the study’s goal but are correlated with the feature of interest. Here, we discuss three approaches: subtracting correlations, residual fitting and feature shuffling. For the first method, two models are created, one with and one without the feature of interest. If the reconstruction or prediction accuracies significantly increase when the feature of interest is included, the feature of interest contributes unique information to the model. This method has been applied in previous studies (e.g. Di Liberto et al., 2015; Brodbeck et al., 2018a; Gillis et al., 2021b). The disadvantage of this method is that it is too convervative. Only the unique contribution of the feature of interest compared to the control features is captured. As shown on Figure 3, some control features, like the envelope, can contain information which is also captured by word surprisal. Therefore, the linguistic information common to both features will be attributed to the envelope. Another disadvantage is that the degrees of freedom change between the two models. This is especially important to consider when the cross-validation technique is not applied. If this approach is used with ridge regression, Crosse et al. (2021) suggests using banded-ridge regression as different features might require different regularisation.

Another approach is to look at the model’s residuals with the control features. In this case, first, a model is created using the control features. Subsequently, the predicted EEG or reconstructed envelope is subtracted from the actual signal, creating the models’ residuals. Then a new model is created using the features of interest and these residuals. If a significant prediction or reconstruction accuracy is achieved, the feature of interest explains variance in the EEG responses, which is not explained by the control features. Similar to the subtracting correlation approach, this method is very conservative as it only considers the unique contribution of the model.

The last approach is shuffling of the features. This approach is used in studies by Broderick et al. (2018, 2021). Now two models include the control features combined with either the feature of interest or a shuffled version of the feature of interest. If the model with the feature of interest performs significantly better than the model with its shuffled version, the feature contributes significantly to the model. Note that shuffling of the feature should be done with care. For impulsive features, shuffling the feature can be done by preserving the timing of the pulses, but changing the amplitudes. For continuous features, the shuffled feature can be created by filtering noise with the same spectrum as the feature of interest. In this method, the number of features is kept constant to investigate the feature’s added value. However, a disadvantage is that the inter-feature correlation is not preserved. For example, returning to Figure 3, if the feature of interest is word surprisal, the correlation between word surprisal and envelope gets lost, i.e., the envelope captures no effect of the shuffled word surprisal. The loss of the inter-feature correlation might affect the model performance, mainly if the cross-validation technique is not applied.

Although each approach has disadvantages, they are all valid and used in different studies. However, good control conditions should always be considered to evaluate whether or not a feature captures the desired effect. For example, for a linguistic feature, this might be a foreign language, vocoded speech or time reversed speech. However, each of these control conditions also has its drawbacks. Depending on the choice of a foreign language, the speaker and language structure can vary, which affects neural tracking. Although vocoded speech can preserve the speech envelope, other speech cues are lost. Time reversed speech has a limitation that the onsets become unnatural, i.e. acoustic boundaries occur at the end of the sound instead of the beginning. An overarching disadvantage is that the listener’s attention may drift over the course of the control stimulus. As the stimulus is not understandable, it is more challenging to listen attentively to the stimulus.

## 4. Neural tracking of the f0

Neural tracking of the fundamental frequency of the voice, or f0-tracking, is used to investigate how the f0 is represented in the brain activity (Forte et al., 2017; Etard et al., 2019; Van Canneyt et al., 2021c). The f0 is a periodic modulation in the speech signal generated by vocal fold vibration during speech production. It is related to the perception of pitch. The f0 of adult speakers typically ranges from 85 to 300 Hz, with male and female voices situated respectively at the lower and higher ends of the range. The f0 is an essential characteristic of the human voice, and it is vital to convey intonation and emotion. However, proper perception of the f0 is not required for speech intelligibility (e.g. cochlear implant listeners). Nevertheless, f0-tracking can provide information on the quality of fine temporal processing in the early stages of the auditory pathway, which is the foundation for proper speech processing in the brain.

Temporal processing of the f0 in the human auditory system happens through the synchronisation of the activity of the neurons to the f0 modulations, i.e. phase-locking. Due to the relatively high frequency of the f0 modulations, this phase-locking occurs mainly in peripheral and subcortical stages of the auditory pathway, up to the upper brainstem. Neurons at cortical stages have poor phase-locking above 100 Hz and are therefore less likely to contribute to f0tracking (Joris et al., 2004). However, it has been shown that early cortical contributions to f0-tracking responses (and FFRs) can occur for low-frequency stimuli (85-100 Hz, e.g. low male voices) (Coffey et al., 2016, 2017; Van Canneyt et al., 2021c).

F0 tracking analysis requires an f0 feature that represents the f0 modulations in the presented speech. The f0 feature can be extracted from the speech stimulus in various ways. A simple yet effective way is to band-pass filter the stimulus in the range of the f0 (Etard et al., 2019; Van Canneyt et al., 2021c). An example of this type of feature is provided in panel A of figure 2. More complicated and computationally expensive techniques have also been explored, including empirical mode decomposition (Etard et al., 2019; Forte et al., 2017) and auditory modelling (Van Canneyt et al., 2021b). Constructing an f0 feature that approximates the expected neural response using auditory modelling has proven particularly effective, nearly doubling the reconstruction accuracies obtained with the neural tracking analysis (Van Canneyt et al., 2021b). This is likely explained by the fact that the auditory model is more physiologically valid. Moreover, the auditory model simulates the contribution of the higher harmonics to the f0 response. Because the neural response is also driven by the harmonics and not just by the f0, the level of measured f0 tracking will increase.

**Figure 2:**
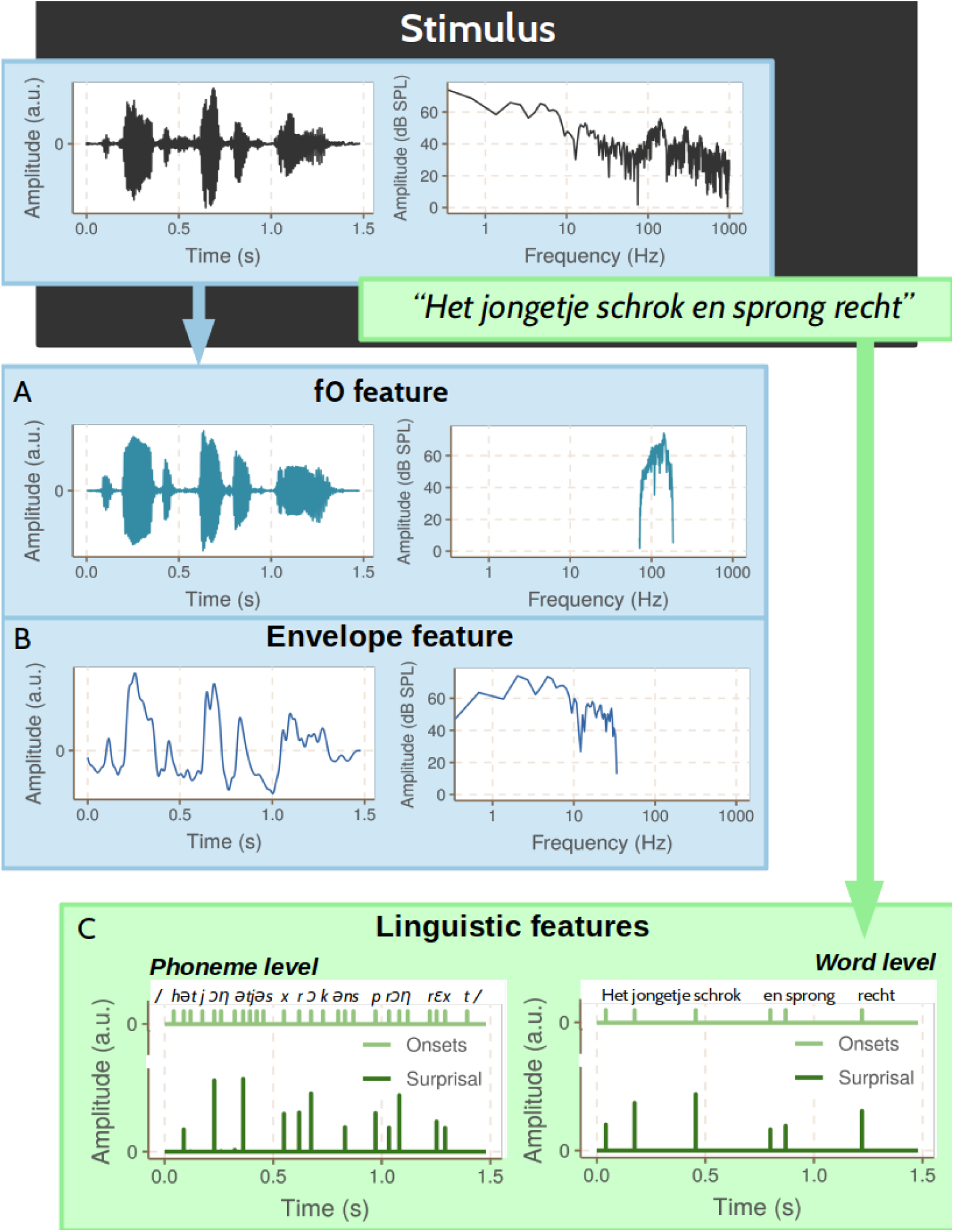
Example of stimulus and derived features for an example sentence by a male speaker. The f0 (panel A) and envelope feature (panel B) are derived from the stimulus waveform, whereas linguistic features (panel C) are derived from the stimulus transcription. The f0 and envelope features concern different spectral ranges, with the envelope focusing on low frequencies (< 50 Hz) and the f0 focusing on higher frequencies (∼ 85 − 300 Hz). Linguistic features can focus on different segmentation levels, including phoneme level and word level. Panel C visualises an example onset and surprisal feature for each level.

Section 1 of figure 4 shows the results of a typical encoding modelling analysis for f0-tracking. These results were obtained by filtering the EEG responses between 75 and 175 Hz and referencing to the average EEG response (32 participants; Van Canneyt et al. (more details regarding the preprocessing are described in 2021c)). The data set used for this visualisation (and all others in figure 4) contained 64-channel EEG data from 32 young normal-hearing subjects measured in response to male-narrated speech (Accou et al. (dataset from 2021)). Panel A shows the mean TRF across subjects for the channel selection indicated in pink on panel B. The TRF for each subject is plotted as well to indicate the variance. The TRFs in this example are absolute value of the TRFs of the stimulus and the Hilbert transformed EEG. This technique aids with interpretation as the brainstem response may occur at a phase that is different from that of the f0 (for more information, see Van Canneyt et al. (2021c); Forte et al. (2017)). The TRF pattern indicates that the activity in the auditory system (∼ EEG) best reflects the f0 information (∼ the feature) at a latency of about 10-25 ms. Panel B of figure 4 presents an example f0 tracking topographic map with common-average rereferencing at 15 ms latency. The topographic map indicates strong response activity in the centre of the scalp and across the back of the head. The temporal and spatial response patterns are consistent with dominant f0related activity in the upper brainstem and early cortical regions. Saiz-Alía and Reichenbach (2020) has performed detailed computational modelling of the subcortical sources of the f0 tracking response, demonstrating important contributions from the cochlear nuclei and the inferior colliculus. Van Canneyt et al. (2021c) argues for additional contributions from the right primary auditory cortex for f0 tracking of low-frequency voices.

**Figure 3:**
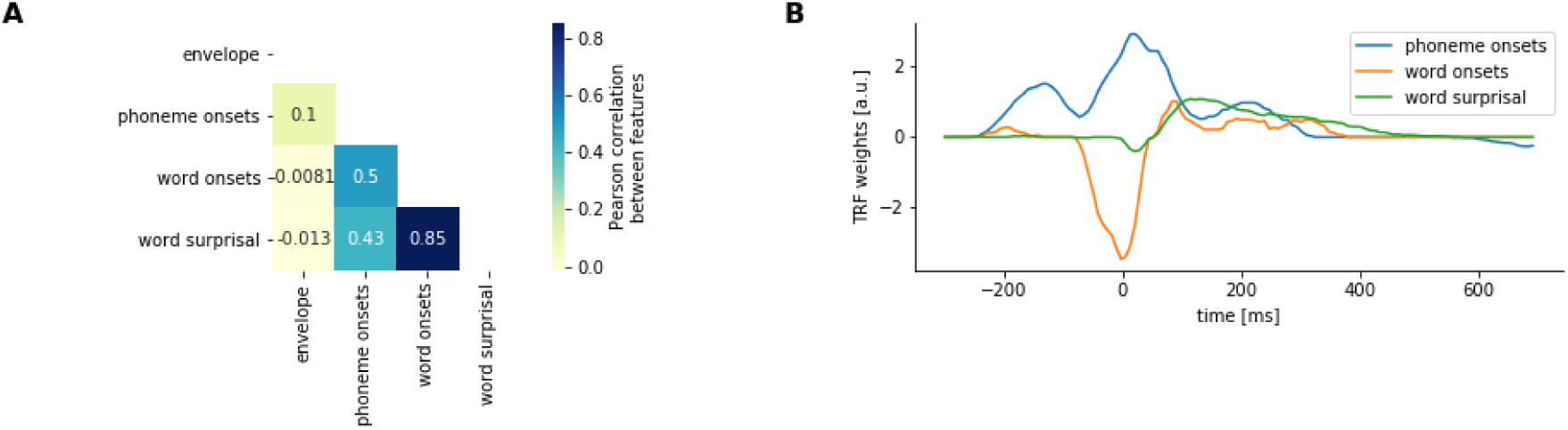
Illustrative example of the inter-feature correlation for the stimulus used in Figure 2. Panel A shows the correlation between the different features. Panel B shows the TRF for phoneme onsets, word onsets and word surprisal when trying to predict the envelope of the stimulus.

**Figure 4:**
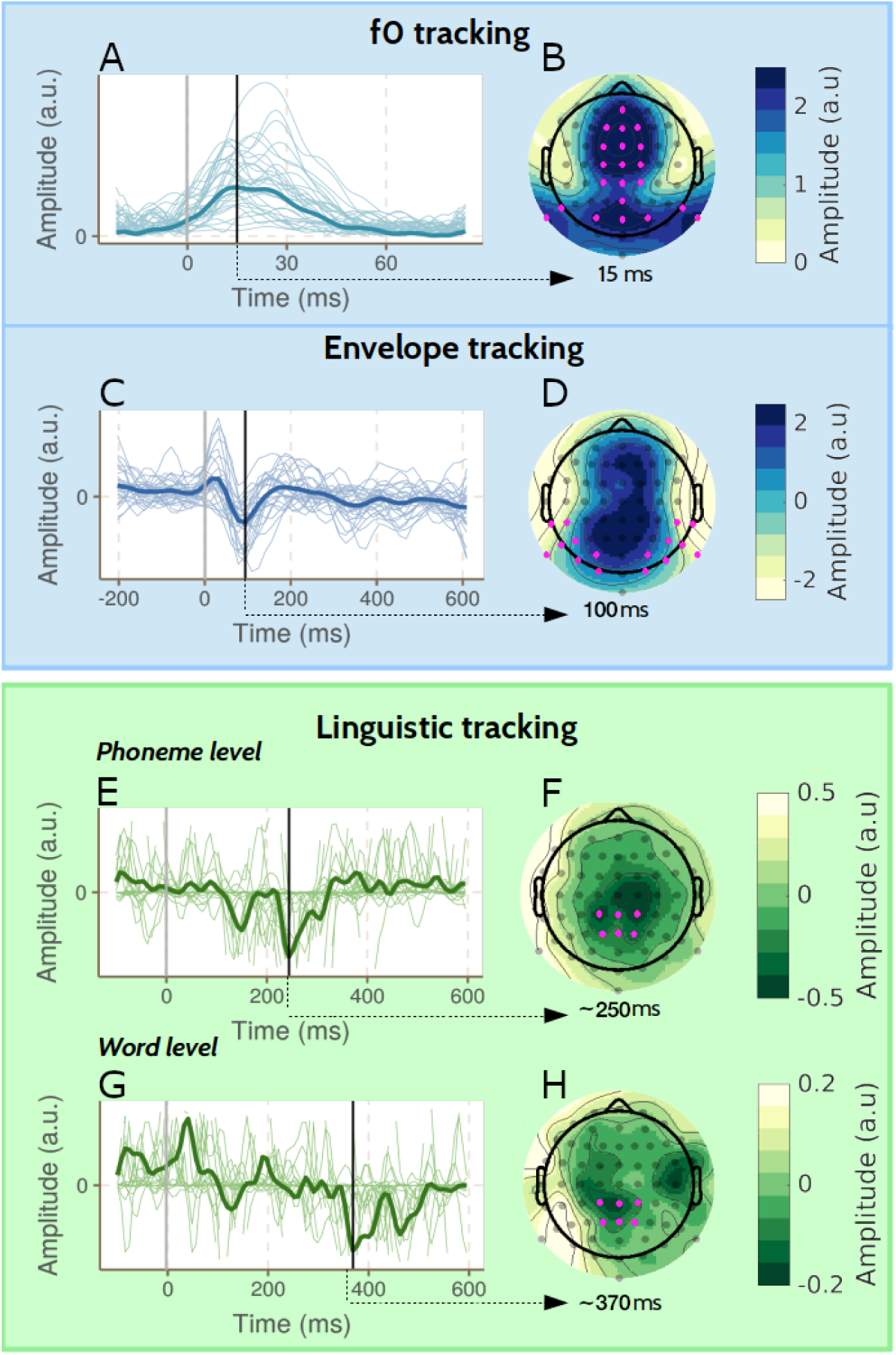
Example of encoding modelling results: TRFs and topographic maps. The figure is divided into three sections on f0-tracking, envelope tracking and linguistic tracking, respectively. For each type of tracking, an example mean TRF (+ individual TRFs) is presented (panel A, C, E and G), together with a corresponding topographic map at an important latency (panel B, D, F and H). These TRFs are estimated for the same participants and the same speech material of around 15 minutes long. The channels indicated with pink on the topographic map represent the channel selection used to obtain the corresponding TRF. Note the drastically different time scales in the TRFs, reflecting the presence of neural activity at different latencies for each feature. In panel A, the TRFs are visualized of the stimulus and the Hilbert transformed EEG, similarly to Van Canneyt et al. (2021c) and Forte et al. (2017).

Although f0-tracking was only recently developed, the technique has led to several interesting findings. Forte et al. (2017) and Etard et al. (2019) have demonstrated that the f0 tracking response holds information on selective attention, possibly indicating that neural mechanisms for attention influence the brainstem. Kulasingham et al. (2020) and Van Canneyt et al. (2021a) have investigated how the age of the listener impacts f0 tracking. Kulasingham et al. (2020) found no age effects using MEG, which is most sensitive to cortical sources. In contrast, Van Canneyt et al. (2021a) found a significant reduction in response strength with advancing age using EEG (which is more sensitive to subcortical sources than MEG and will capture both cortical and subcortical sources). This observation is in line with an age-related decrease in the phase-locking ability of the subcortical (and early cortical) auditory system. Van Canneyt et al. (2021a) also studied the effect of hearing loss and found increased f0-tracking responses in participants with hearing impairment compared to age-matched controls. The response enhancement was due to additional cortical activity phase-locked to the f0 (with a latency of ∼40 ms), likely compensating for the reduced quality of bottom-up auditory input due to diminished peripheral auditory sensitivity. Moreover, the amount of additional compensatory cortical activity was significantly related to the pure tone average (PTA) hearing loss of the participant. As such, a significant relationship exists between the degree of hearing loss of an individual and the strength of their f0 tracking response.

At the moment, f0-tracking also has some limitations, which future advances may mitigate. One of the main issues is auto-correlative smearing in TRFs and topographic maps because the f0 stays relatively steady over multiple f0 periods. This periodic smearing over latencies can be somewhat mitigated with Hilbert-transformed TRFs, which disregard phase information. However, TRF and topographic map interpretation are still limited to the most dominant peaks (see Van Canneyt et al. (2021c) for more details). A second limitation is that the f0 is only present in speech during voiced sounds (∼ 50-60 % of the time) and not during unvoiced sounds (∼ 40 % of the time), including silences. During analysis, these unvoiced sections in the speech stimulus (and corresponding sections in the EEG) are disregarded. As a result, only about half of the measured data can be used to analyse f0-tracking, increasing the required measurement time. Another limitation is that the f0 tracking response is reduced for voices with higher and more variable f0, leading to weak and often non-significant responses for typical female voices. This occurs because neural phase-locking ability is decreased for higher and more variable f0s, especially for cortical sources. As such, the stimulus choice has a large impact on the f0 tracking response. A final limitation is that f0-tracking requires careful interpretation: f0-tracking reflects the capability of the auditory system to phase-lock to the f0, but it does not reflect the ability of a person to perceive pitch or speech in general. Neural tracking analyses with other features help complete the picture. Lastly, the use of the f0 feature is still in its infancy with only a limited number of published studies. Nevertheless, it seems to be a promising approach to evaluate neural responses at the level of the brainstem.

## 5. Neural tracking of the speech envelope

The speech envelope consists of slow-varying modulations (< 50 Hz) in the speech signal. It contains acoustic temporal information (Rosen, 1992) but also reflects phonemes, syllables and word transitions (Peelle and Davis, 2012). Moreover, it also correlates with the area of the mouth opening during articulation (Chandrasekaran et al., 2009). Therefore it is not surprising that research indicates that the envelope is an essential acoustic cue for speech intelligibility (Shannon et al., 1995; Drullman et al., 1994a,b).

Measuring envelope tracking can be used to analyse the neural encoding of the speech envelope during speech perception (Ding and Simon, 2012a; O’Sullivan et al., 2015; Vanthornhout et al., 2018). From animal studies (Wang et al., 2008) and human studies with electrocochleography (ECoG), it is known that the speech envelope is processed in the primary auditory cortex, specifically in Heschl’s Gyrus (Nourski et al., 2009). A growing body of evidence demonstrates that envelope tracking is a requirement for speech understanding. Multiple studies show that neural tracking of the speech envelope is strongly correlated with behaviourally measured speech intelligibility (e.g. Ding et al. (2014); Vanthornhout et al. (2018); Lesenfants et al. (2019); Iotzov and Parra (2019); Verschueren et al. (2021)). As a specific example, Vanthornhout et al. (2018) found a significant correlation of 0.69 between the speech reception threshold (SRT) estimated based on envelope tracking and the SRT measured with behavioural speech audiometry.

Although the broadband envelope can be used, it is also possible to study the neural tracking of specific frequency bands of the envelope. Envelope tracking responses are most commonly investigated in the delta band (0.5-4 Hz), theta band (4-8 Hz) and gamma band (> 30 Hz) (Ding and Simon, 2013; Verschueren et al., 2021; Molinaro and Lizarazu, 2017). The lower envelope frequencies are often the main interest as they correspond with word onsets and the syllabic rate of the speech, which is hypothesised to be crucial for speech intelligibility. Higher envelope frequencies are typically related to the onsets of phonemes. Some studies suggest that speech intelligibility, i.e., how well a person can understand speech, is specifically related to the theta band (4-8 Hz) and not the delta band (1-4 Hz) (Ding and Simon, 2013; Peelle et al., 2013). Other studies indicate the opposite (Verschueren et al., 2021; Molinaro and Lizarazu, 2017; Di Liberto et al., 2018). In our opinion, the outcome may depend on the speech material. The syllabic rate is often very close to 4 Hz, so envelope tracking to a slow speaker could be more dominant in the delta band, while envelope tracking to a fast speaker could be more dominant in the theta band.

Envelope tracking responses can be analysed using a decoding or encoding model, or using a strategy that combines a decoding and encoding approach in one model. In any case, the model requires an envelope feature extracted from the stimulus waveform. In essence, the envelope reflects the modulation of the signal and can easily be obtained by taking the absolute value of the Hilbert transform. Although this is a prevalent method, it is not the best choice as it disregards auditory processing. Two important aspects of auditory processing need to be taken into account to better approximate human envelope processing. First, the stimulus should be split into frequency bands before the actual envelope extraction process to mimic how the basilar membrane in the cochlea divides a sound stimulus into different auditory filters. Second, the compression and non-linear behaviour of the auditory system should be accounted for. To incorporate these factors in the envelope extraction process, complex computational models of the auditory periphery can be used (Yang et al., 2015; Bruce et al., 2018). However, Biesmans et al. (2017) evaluated various extraction methods in an auditory attention decoding (AAD) paradigm and proposed a simplified approach. They found that a combination of a gammatone filterbank, which simulates the auditory filters on the basilar membrane, followed by a power law to account for compression and non-linearity in the auditory system, performed equally well as the more complex and computationally expensive auditory models. An example envelope feature obtained using this technique is provided in panel B of figure 2.

The results of a typical encoding modelling analysis using ridge regression for envelope tracking are visualised in section 2 of figure 4. These results were obtained by highpass filtering the EEG responses above 0.5 Hz and referencing to the average EEG response (32 participants; Vanthornhout et al. (more details regarding the preprocessing are described in 2019)). Panel C presents the mean TRF, averaged over subjects and a channel selection (indicated in pink on panel D). The TRFs of the individual subjects are visualised with a thin line to indicate the variance. The TRF displays three distinct peaks. The P1 peak (50 ms), the N1 peak (93 ms) and the P2 peak (170 ms). This typical P1-N1-P2 complex is also found in AEP studies with impulse-like stimuli and can thus be used to infer the neural source of the peaks. The P1 peak originates in Heschl’s Gyrus, and the N1 peak originates in the Superior Temporal Gyrus (O’Sullivan et al., 2019b; Steinschneider et al., 2011). The origin of the P2 peak is less clear but is probably in the (higher) auditory cortex (Godey et al., 2001). The topographic map shows negative weights for the temporal channels and positive weights for the central channels. This distribution is an indication of a dipole located near the auditory cortex. Without analyses in source space, the exact location is difficult to pinpoint.

Over the past decade, envelope tracking has been used to study, among others, how cortical speech processing is affected by individual factors like age and hearing status. Decruy et al. (2019) and Brodbeck et al. (2018b) found stronger envelope tracking for older participants compared to younger participants, even though older adults typically have more difficulty understanding speech. Similarly, Decruy et al. (2020b) and Fuglsang et al. (2020) found increased envelope tracking for hearing-impaired listeners compared to age-matched normal-hearing listeners. The enhanced tracking in older listeners or listeners with a hearing impairment may be explained by a compensatory central gain mechanism (Parthasarathy et al., 2019; De Villers-Sidani et al., 2010; Chambers et al., 2016), recruitment of additional cortical resources (Brodbeck et al., 2018b; Gillis et al., 2021a) and increased listening effort and attention (Decruy et al., 2020a; Vanthornhout et al., 2019; Lesenfants and Francart, 2020). This shows that it is also important to conduct subject-specific analyses and not only at group-level measures. With an innovative artefact removal technique, Somers et al. (2019) succeeded to analyse envelope tracking for cochlear implant listeners as well. For both hearing-impaired listeners (with simulated amplification) (Decruy et al., 2020b) and cochlear implant listeners (Verschueren et al., 2019) the tracking strength was significantly correlated to behaviourally-measured speech intelligibility, indicating a similar relation with speech intelligibility as observed for normal hearing listeners (Vanthornhout et al., 2018).

One challenge with envelope tracking is that its functional interpretation is unclear. The main complicating factor is that the envelope is highly correlated with linguistic cues, like the onsets of words and syllables. As such, the envelope represents multiple unique features that all may contribute to the observed neural tracking response and are hard to disentangle. In addition, the interpretation of envelope tracking is complicated because it is modulated by top-down effects, such as attention and audio-visual integration (O’Sullivan et al., 2019a). A final challenge is that the exact relation between envelope tracking and speech intelligibility remains a point of discussion (Ding and Simon, 2014; Brodbeck and Simon, 2020). Multiple studies have shown that the level of envelope tracking reflects experimental changes in speech intelligibility (Vanthornhout et al., 2018; Lesenfants et al., 2019; Verschueren et al., 2021), even in the case of degraded speech with an intact envelope (Ding et al., 2014). However, it is unlikely that envelope tracking is a direct reflection of successful speech intelligibility as neural tracking responses have been observed for non-speech signals (Zuk et al., 2021) and foreign languages (Etard and Reichenbach, 2019). As such, envelope tracking is likely necessary but not sufficient for speech intelligibility. Linguistic features can be used to gain further insight into how the brain processes the meaning of speech, i.e. speech intelligibility.

## 6. Neural tracking of linguistic features

Recent studies focus on linguistic speech features in pursuit of an accurate neural marker of speech intelligibility. While the f0 and speech envelope are derived from the acoustic waveform of the speech, linguistic features are derived from the content of the speech. Proper encoding of these features in the brain requires accurate linguistic and not mere acoustic processing.

Linguistic features can be divided into two categories. Features in the first category denote lexical onsets. They represent (aspects of) a sequence of small building blocks that make up spoken language, e.g., sequences of phonemes, phonetic features, words, or specific word categories like content and function words (Di Liberto et al., 2015; Lesenfants et al., 2019). These features are sparse arrays consisting of zeros with a fixed, non-zero entry (∼ impulse) at the onset of each lexical building block (see features in light green on Panel C of figure 2). The neural responses to lexical onset features are not straightforward to interpret. As phonemes, syllables, and words coincide with acoustic cues, the associated neural response is neither purely lexical nor acoustic.

Features in the second category reflect higher-level linguistic aspects of the speech, e.g., how familiar, predictable or surprising a word or phoneme is in its context. These features can be applied on three levels, which require different amounts of linguistic context: (1) at the level of a phoneme (e.g., phoneme surprisal or cohort entropy (Di Liberto et al., 2019; Brodbeck et al., 2018a)), (2) at the level of a word (e.g., word frequency or word surprisal (Weissbart et al., 2019; Koskinen et al., 2020)), and (3) at a semantic contextual level (e.g., semantic dissimilarity (Broderick et al., 2018)). These features are sparse arrays, similar to lexical onset features. However, in this case, the impulse amplitude at each onset is not fixed but modulated by the linguistic information of the specific phoneme or word (see features in dark green on Panel C of figure 2).

The fact that linguistic features are sparse arrays consisting of mostly zeroes with some non-zero entries (∼ impulses), makes them different from the continuous f0 and envelope features and poses challenges for response analysis. In decoding modelling, the reconstructed feature is compared to the actual feature. However, sparse features are challenging to reconstruct from a continuous signal (the EEG), as we can only reconstruct a continuous signal from the continuous input using a linear model. This problem does not occur for encoding modelling, where the non-sparse predicted EEG is compared to the actual EEG. Therefore the encoding model is a more common choice for analysis with linguistic features.

Panels E-H of figure 4 present a visualisation of the results of a typical encoding modelling analysis for linguistic tracking with phoneme suprisal and word suprisal features. These results were obtained by filtering the EEG responses between 0.5 and 25 Hz and referencing to the average EEG response (32 participants; Gillis et al. (more details regarding the preprocessing are described in 2021b)). The TRFs at both phoneme (panel E) and word level (panel G) show a negative response, situated centrally in the topography (panel F and H), around respectively 250 and 350 ms. The earlier response peak for phonemes compared to words is consistent with the hierarchy of the language processing of these linguistic building blocks, i.e., the phonemes making up a word are processed before the word’s surprisal can be estimated. Moreover, the response to word surprisal resembles the N400 response, which is classically observed in ERP paradigms (Lau et al., 2008). These congruent topographic responses indicate that this small and specific language response can also be observed when listening to natural running speech rather than stand-alone sentences.

Measuring neural tracking of linguistic features is an exciting avenue to test psycho-linguistic theories of speech understanding. It is accepted that listeners use linguistic context to continuously adapt expectations of upcoming concepts, words and phonemes, However, how these expectations are integrated with what is actually being perceived is unclear. Brodbeck et al. (2021a) showed that the neural prediction of an upcoming phoneme or word relies on contextual processing in a parallel manner, combining both bottom-up and top-down processing. Additional evidence of the presence of top-down processing comes from Heilbron et al. (2020) who observed that higher-level predictions influence the predictions at lower levels (i.e., word prediction affects the predictions at the phoneme level).

Additionally, linguistic features allow investigating to what extent speech is understood given the language proficiency. Di Liberto et al. (2021) investigated neural tracking in Mandarin speakers with different levels of English proficiency. Interestingly, the magnitude of central negative response to semantic dissimilarity around 400 ms increased with proficiency.

Another exciting research path is the disentanglement of acoustic and linguistic neural processing. Verschueren et al. (2022) disentangled acoustic and linguistic neural processing by changing the speech rate, which kept the linguistic content the same while varying the acoustic properties and the intelligibility of the speech. As the speech rate increased, the neural tracking of acoustic properties increased. This means that better time-locking was observed, even though the speech became harder to understand. In contrast, neural tracking of linguistic properties decreased with increasing speech rate. This indicates that linguistic tracking provides a more accurate objective measure of speech intelligibility.

The two studies mentioned above indicate that neural tracking of linguistic features encodes aspects of neural language processing. These findings open doors to study language development in young children or to objectively determine speech understanding.

Linguistic speech features can also provide insight into age-related speech intelligibility deficits. We are aware of two studies investigating speech intelligibility deficits in older adults. Although Mesik et al. (2021) did not report differences, Broderick et al. (2021) reported that older adults rely less on semantic features than younger adults. Furthermore, they showed that older adults who relied more on this semantic mechanism showed higher verbal fluency than older adults with weaker semantic tracking.

Linguistic tracking is an up-and-coming research technique but has a few difficulties. Firstly, the inter-feature correlation should be taken into account. The linguistic features coincide with the boundaries of phonemes and words. These boundaries are often associated with high acoustic power; therefore, it is necessary to carefully control for acoustic properties of the speech when evaluating linguistic features. If not, the speech tracking analysis might be biased to find spurious significant linguistic features due to its correlations with acoustic features (Daube et al., 2019). We proposed some approaches to deal with this inter-feature correlation in the section 3.1.

Secondly, the analyses are often based on encoding modelling due to sparse features. Prediction accuracies, i.e. correlations, obtained with encoding models are typically small in magnitude: only around 3 to 7% of the variance in the EEG signal can be explained by neural responses time-locked to the presented stimulus. Moreover, most of this variance is explained by acoustic characteristics of the speech, as these lower-level acoustic features evoke responses over large parts of the auditory system. In contrast, linguistic tracking targets the neural response from a precisely localised neural process related to intelligibility. Therefore, the associated magnitudes of these neural processes measured at the scalp level are much smaller. As the prediction accuracies of the encoding model are small in magnitude, finding a significant improvement of the linguistic feature over and beyond acoustic features requires enough observations (e.g. an improvement of ∼1% corresponds to an increase in prediction accuracy of 3.4 × 10^−4^ approach as described above (Gillis et al., 2021b)).

Thirdly, estimating the TRF to these linguistic features requires a lot of data. In Figure 1, a 15-minute story is used to estimate the model, which is substantially shorter than most studies which evaluate linguistic tracking of speech in 45 to 60 minutes. This difference in the amount of data explains why the TRF pattern is noisier. The more data is used to estimate the model, the better the brain response to linguistic tracking is characterised, and the more prominent the peaks are. A method to overcome this is to estimate a subject-independent model whereby the data of different subjects is combined to ensure enough data.

Another difficulty is that current studies investigating the effect of linguistic neural tracking only evaluate population differences, such as comparing young to old or comparing a baseline to a complete model. As the effect is small and the TRFs show a high between- and intra-subject variability, it is challenging to extract an objective measure at a subject-specific level. Future research should focus on making the TRF patterns and prediction accuracies more reliable and robust at a subject-specific level.

## 7. Caveats

A first caveat of the studies investigating neural tracking is that most studies focus on models assuming a linear relationship between speech and EEG responses. However, whether or not this assumption is valid remains unanswered. Additionally, there is no ground truth of which speech features the brain tracks. Therefore, investigating which speech features are tracked by the brain remains an explorative search which might lead to suboptimal results.

A second caveat is the comparison of neural tracking of speech with behavioural measures of speech understanding. Behavioural measures can be obtained by, for example, sentence recall tests. However, when using continuous speech, these tests are not suitable because longer text fragments cannot be recalled. As a result, the speaker of the behavioural measures is often different from the speaker of the stimuli used for the neural measurements. This can be problematic as speaker characteristics can affect the neural tracking of speech. As sentence recall tests are unsuitable, the current field of research lacks a good evaluation of speech understanding for continuous text. Currently used metrics are content questions or subjectively rated speech understanding, i.e., the participant’s answer to the question ‘how much did you understand?’. To overcome the intersubject variability of these subjective ratings, the self-assessed Békesy procedure can be applied to rescale the subjective ratings towards a more objective alternative (Decruy et al., 2018). However, these metrics remain sub-optimal as content questions rely heavily on attention effects and subjectively rated speech understanding introduces a bias.

Another caveat is that neural tracking is sometimes confused with neural entrainment. Both concepts underlie different assumptions. Neural tracking assumes that the neural responses time-lock to different sound features. Speech features evoke a cascade of different responses, such as responses that are time-locked to the acoustical, lexical, and linguistic features. In contrast, neural entrainment assumes that the neural responses phase-lock to the stimulus. This theory assumes that oscillations are present in the brain without stimulation. When a rhythmic sound, such as speech, is heard, these oscillations reset their phases and become synchronised with the dominant phase of the speech signal (Peelle and Davis, 2012). Phase-locking can be measured, for example, using inter-trial correlations (Ding et al., 2014). As a result, neural entrainment involves more assumptions than neural tracking. Therefore, it is possible that speech understanding can occur when there is no neural entrainment, for example, when the speech signal is very arrhythmic (Peelle et al., 2013). On the other hand, having neural tracking does not guarantee that the speech signal was understood correctly.

## 8. Clinical applications of measurements of neural tracking responses

To provide care to people with hearing problems, it is useful to review the merits and limitations of all (objective) audiological measures and investigate how the measures may be combined to form a complete assessment of the auditory system.

The current gold standard methods, i.e. tone and speech audiometry, have proven their worth. However, they are challenging in crucial populations like young children or people with a cognitive impairment like dementia. Objective measures for sound perception like the ABR and the ASSR have been introduced in the clinical toolset to remedy this. However, there is no clinically available objective measure of speech intelligibility. Since speech intelligibility is the basis for human communication, this is a significant gap to fill. Various populations may benefit from such a measure, including young children, stroke patients (especially those with aphasia) and people with dementia.

Measurement of neural tracking is a versatile tool as the amount of tracking to the different speech features and thus in different parts of the auditory system can be measured. Based on a single twenty-minute long EEG recording, a wide range of speech processing abilities may be assessed simultaneously (incl. f0 tracking, envelope tracking, phonetic processing, phonemic processing, syntactical processing and even linguistic and emotional processing). This versatility may lead to an objective assessment of both auditory and language abilities. Moreover, measuring neural tracking is easily automated, paving the way to improved automated screening, diagnostics, and automatic fitting of auditory prostheses, or even auditory prostheses that continuously adapt themselves to the listener based on their brain activity (Geirnaert et al., 2021).

Future studies preparing for clinical implementation may need to shift focus from group-level analyses towards subject-specific analyses. Going towards a subject-specific analysis is key to allow its feasibility as a clinical marker. Although the magnitude of neural tracking might be intrinsically different between different subjects, relative differences between different conditions can be used as a diagnostic marker. Additionally, future studies may focus on which combination of stimulus features provides the most information and how these can be optimally analysed. As the features are highly correlated, special care needs to be taken to investigate the effect of each feature (Gillis et al., 2021b). Subsequent research efforts are also required to decide on the best speech stimuli (required to work well for all types of tracking) and the best EEG measurement set-up, including the number of EEG electrodes and their position (Montoya-Martínez et al., 2021) to reduce recording time. Furthermore, it is important to set best practices of how the measures of neural tracking can be used (Crosse et al., 2021) and to have insight into how the preprocessing influences the results (de Cheveigné and Nelken, 2019). It is also essential to validate the measures in a comprehensive sample of the population, including participants of all ages and with various audiological and non-audiological pathologies (for a review, see Palana et al. (2021)). Finally, the neural tracking results need to be transformed into an easy-to-interpret set of scores and visualisations to allow for intuitive use by clinicians.

## 9. Acknowledgements

The authors would like to thank Bernd Accou, Wendy Verheijen and their students for collecting the dataset used for the examples in this article. The research received funding from the European Research Council under the European Unions Horizon 2020 research and innovation programme (grant agreement No. 637424, ERC starting grant to Tom Francart). Jana Van Canneyt and Marlies Gillis are both supported by a PhD grant for Strategic Basic research by the Research Foundation Flanders (FWO): project number 1S83618N and project number 1SA0620N, respectively. Jonas Vanthornhout is funded by a postdoctoral grant from FWO, project number 1290821N.

